# The Adaptive Brain Across Development: Age Related Changes in Cortical Adaptation

**DOI:** 10.64898/2026.02.23.707381

**Authors:** Carmel Moalem, Ofri Levinson, Sagi Jaffe-Dax

**Affiliations:** Sagol School of Neuroscience, Tel Aviv University; School of Social Science, Tel Aviv University

**Keywords:** fNIRS, cortical adaptation, cortical development, functional connectivity

## Abstract

How does the functionality of the cortex change from infancy to adulthood to support the developmental cognitive shift from learners to performers? Cortical adaptation is a simple neural mechanism which plays a key role in learning and memory encoding, but little is known about how it develops across the lifespan. Both infants and adults have been found to respond differently to repeating audio and visual stimuli, suggesting differences in cortical adaptation throughout development. However, studies typically approach these populations through different paradigms and interpret the results in terms of different cognitive models. To overcome these issues, we implemented an identical paradigm across all age groups to examine cortical adaptation and its developmental trajectory. We used functional near infra-red spectroscopy (fNIRS) to chart how different regions in the infant, child and adult brain respond to repeating audiovisual stimuli at varying inter-stimulus intervals (ISIs), using cortical adaptation as a proxy for implicit memory dynamics. We found faster recovery from adaptation in infants compared to children and adults. Specifically, there was an interaction between stimulus presentation rate and age in the right temporal, left parietal and occipital cortical areas. There was also a developmental progression in functional connectivity, with infants displaying significantly lower correlations between regions of interest than children and adults. Taken together, we suggest these findings may reflect the developmental trajectory of cortical adaptation from a learning system optimized for maximal information intake and minimal filtering of stimuli to a specialized integrative system that efficiently filters and adapts to information.

**Highlights:**

- Cortical adaptation is a fundamental mechanism involved in memory and learning, but not much is known about how it develops throughout the lifespan.
- An identical fNIRS paradigm across 3 different age groups reveals significant differences in cortical adaptation between infants, children and adults.
- Functional connectivity revealed foundational connections present from infancy, growing stronger and into a specialized adaptation system with age.

These findings suggest a developmental transition from a system optimized for maximal information intake to a specialized learning system, capable of filtering redundant information.

## 1. Introduction

The human brain undergoes significant structural and functional changes throughout development that shape perception from infancy to adulthood. Cognitive development can be understood as a dynamic interaction between sensory information and cognitive structures.

Piaget’s theory of cognitive development describes this process as an interplay between assimilation and accommodation, where new sensory information is incorporated into existing cognitive structures while also reshaping them (Block, 1982). These overall differences help explain why infants and children sometimes outperform adults on specific perceptual tasks despite (or because of) their more limited cognitive capacities (Gualtieri & Finn, 2022; Meier & Newport, 1990; Plebanek & Sloutsky, 2017). However, perceptual processing strategies also evolve individually during this development. For instance, younger infants initially prefer familiar stimuli but later exhibit a preference for novel stimuli (Raz et al., 2023). Similarly, while infants and children demonstrate greater reliance on immediate sensory information, adults increasingly incorporate prior experience into their perceptual judgments (Jaffe-Dax et al., 2023).

One fundamental mechanism that is frequently explored in studies on perceptual and cognitive development is habituation, the process by which behavioral responses (e.g., gaze, heart rate) decrease with repeated stimulus exposure (Domsch et al., 2010; Horowitz, 1972; Jeffrey & Cohen, 1971). Developmentally, habituation differs across age groups; namely, infants habituate more slowly, recover faster, and require more stimulus exposure than adults (Arichi et al., 2012; Lloyd-Fox et al., 2019), whereas children also show slower habituation than adults (Muenssinger et al., 2013). While most studies on habituation have focused on behavioral measures, understanding the neural mechanisms underlying this phenomenon can provide further insights into how the brain optimizes information processing throughout development. Cortical adaptation might represent a suitable candidate for such a neural mechanism.

Cortical adaptation, also known as repetition suppression, reflects the brain’s increased efficiency in processing a repeating stimulus through a reduced neural excitatory response over time (Grill-Spector et al., 2005). These changes in neuronal excitability are thought to facilitate memory encoding within experiments and throughout development (Fitz et al., 2020). Cortical adaptation has been widely studied and replicated in neuroimaging research (Kobayashi et al., 2014; Lu et al., 1992; Weigelt et al., 2008) and has been found in both infants and adults (Ignatiadis et al., 2024; Lasky, 1997).

Developmental changes in cortical adaptation may be driven by two key mechanisms. . During early childhood, synaptic density in the cerebral cortex peaks before gradually declining to adult levels through the process of *synaptic pruning* (Casey et al., 2000; Huttenlocher & Dabholkar, 1997; Huttenlocher, 1979). Concurrently, *Myelination,* namely, the formation of myelin sheaths around axons and facilitating faster neural transmission, accelerates during early childhood with key pathways maturing by age three (Branson, 2013). These processes refine neural transmission and create more efficient information processing pathways, enabling better cognitive performance and faster recovery from adaptation (Van Der Knaap et al., 1991).

Although synaptic pruning and myelination refine local neural circuits, brain development also involves large-scale changes in *functional connectivity*; i.e., the temporal coordination of activity between spatially distributed neural regions (Fingelkurts et al., 2005). While basic sensorimotor and visual networks demonstrate synchronization from birth (Lin et al., 2008; Liu et al., 2008; Thomason et al., 2013), higher-order networks, such as the default mode network (DMN), mature more gradually through infancy and childhood (Bulgarelli et al., 2020; Gao et al., 2009, 2014, 2017; Kelsey et al., 2021).The maturation of functional connectivity networks represents a critical aspect of brain development, in that basic whereas and continue to strengthen through infancy and childhood. Recent evidence suggests potential interaction between adaptation and functional connectivity in adults (Della-Maggiore & McIntosh, 2004; Kar et al., 2019) however, it is unclear whether and how these processes influence each other during development.

Understanding how cortical adaptation and functional connectivity develop is therefore crucial to uncover the neural mechanisms that support the brain’s increasing capacity for efficient perception and cognition across the lifespan.

When examining adaptation, studies typically fall under one of two approaches: infant studies focus on early learning and memory formation (Oakes, 2010), whereas adult research applies more advanced cognitive models of perception and attention (Chun, 2008b; Raz et al., 2024). Faster adaptation in infancy is linked to advanced cognitive abilities (Oakes, 2010), while atypical cortical adaptation in adolescents and adults has been associated with neurodevelopmental conditions (Gertsovski & Ahissar, 2021; Jaffe-Dax et al., 2017, 2018; Lieder et al., 2019; Perrachione et al., 2016; Turi et al., 2015). These separate research approaches make it difficult to identify the developmental trajectories of cortical adaptation,

### The Present Study

Significant knowledge gaps remain regarding how cortical adaptation and functional connectivity develop and interact across the lifespan. The current study addressed these gaps by simultaneously investigating both phenomena using an identical functional near-infrared spectroscopy (fNIRS) methodology and stimulus protocols in infants, children, and adults. This novel approach bridges traditionally separate research paradigms and makes it possible to isolate fundamental developmental mechanisms. Employing an identical task and neuroimaging modality across all age groups enabled us to map developmental trajectories of cortical adaptation and functional connectivity.

This study investigated two research questions: (1) How do mechanisms of cortical adaptation to repeated stimuli evolve across infancy, childhood, and adulthood? (2) How do patterns of functional connectivity between cortical regions develop across these same stages?

Using fNIRS, we investigated how different cortical regions in the infant, child and adult brains responded to repeating audiovisual stimuli at varying inter-stimulus intervals (ISIs). We hypothesized that infants would exhibit faster recovery than children and adults, with adults demonstrating the slowest recovery. We also hypothesized that functional connectivity would increase progressively with age.

Beyond advancing our understanding of typical neurodevelopment, the findings have implications for the study of atypical development by establishing normative patterns against which deviations associated with neurodevelopmental disorders can be identified

## 2. Method

### 2.1 Participants

The sample was composed of three groups of typically developed individuals: infants (10-14 months; n=41,16 female), children (4.5-5.8 years; n=42, 22 female), and adults (19-33 years; n=44, 16 female).

Several participants were excluded due to suspected developmental disorders (1 infant, 1 child, 1 adult), failure to complete the experiment (3 infants, 2 children), or poor data quality (5 infants, 6 children, 1 adult). The final sample consisted of 31 infants (16 female), 32 children (17 female), and 44 adults (16 female).

Informed consent was obtained from all participants or their guardians. Parents provided consent for the infants, the children provided their assent, and the adults provided their consent.

This study and all the procedures were approved by the Tel Aviv University IRB.

### 2.2 Experimental Paradigm

#### Procedure

Participants were seated comfortably in front of a screen, either alone or with their parents. The children and adults received briefings about the experiment and instructions to remain still while maintaining attention on the screen. Parents of infant participants were also briefed. After obtaining consent/assent, the participants were fitted with a fNIRS cap, engaged in the experiment, and (for children and adults) completed a short memory test. The adults and parents were compensated for their time (approx. $10), and the children and infants also received a t-shirt and a book.

#### Stimulus presentation

While wearing the fNIRS cap the participants were exposed to six randomized trios consisting of audio-visual stimuli (2 seconds), followed by an ISI (2, 3, 4, 6, 8, or 12 seconds). Each sequence repeated eight times within a block (Figure 1). This paradigm is designed to induce adaptation through repeated exposure to identical stimuli, with the assumption that the shorter ISIs will cause a stronger adaptation to the stimuli in all the age groups compared to the longer ISIs. The Inter-Block-intervals (IBIs) lasted approximately 10 seconds, a duration previously shown to be sufficient for infant and child brain responses to return to baseline (Plichta et al., 2007). All experiments were randomized and specific to each participant.

**Figure 1:**
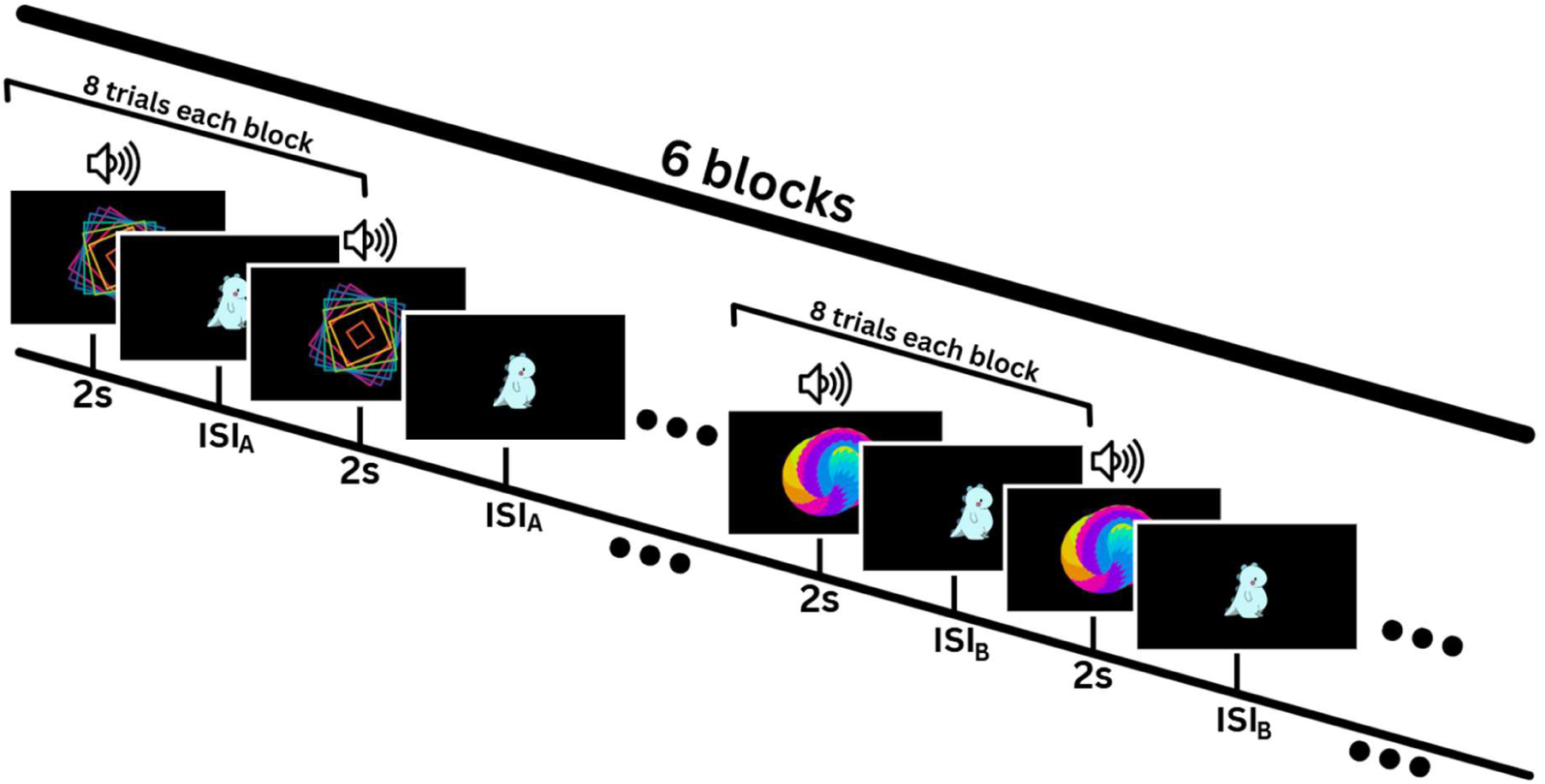
Illustration of the experimental paradigm. An audiovisual stimulus was presented for 2 seconds, followed by an attention getter (ISI) that was 2,3,4,6,8 or 12 seconds long. The stimulus-ISI pairings repeated 8 times per block. Each block included a different stimulus-ISI pairing, with presentation order randomized across participants.

#### Stimulus

Abstract auditory and visual stimuli were used to minimize prior knowledge effects while maintaining engagement through a colorful design.

#### Memory test

Children and adults received instructions to remember the stimuli for a subsequent memory test. This was intended to enhance attention and concentration during the primary experiment.

### 2.3 fNIRS montage and data collection

Functional near-infrared spectroscopy (fNIRS), which measures changes in hemoglobin concentration in the cortex after neuronal activation (Pinti et al., 2018) has emerged as a valuable tool in developmental neuroscience. It is well-suited for studying brain functions in infants and young children (Lloyd-Fox et al., 2009; McDonald & Perdue, 2018; Meek et al., 1998), fNIRS has improved temporal resolution than fMRI, better spatial resolution than EEG, and greater tolerance for movement (Herrmann et al., 2007; Nishiyori, 2016; Quaresima & Ferrari, 2019).

Auditory and visual stimuli have been found to reliably evoke positive fNIRS responses in the temporal and occipital areas, respectively (Harrison & Hartley, 2019; Pollonini et al., 2013; Wijeakumar et al., 2012), consistent with other neuroimaging methods (Cui et al., 2010; Noah et al., 2015), and also have been found to evoke adaptation in similar settings (Emberson et al., 2015). Due to these reasons, we decided to use fNIRS for this research.

A NIRx fNIRS system (NIRSport2, NIRx) was employed for data collection. The montage, designed using the NIRx NIRSITE application, consisted 16 sources and 15 detectors, resulting in a total of 42 channels (Figure 2). The adult montage also included eight additional short channels - four for each hemisphere - to remove superficial physiological noise. The montage covered the bilateral frontal, temporal, parietal and occipital cortical areas. For all age groups channel spacers were used to preserve equal distances between the optodes.

**Figure 2:**
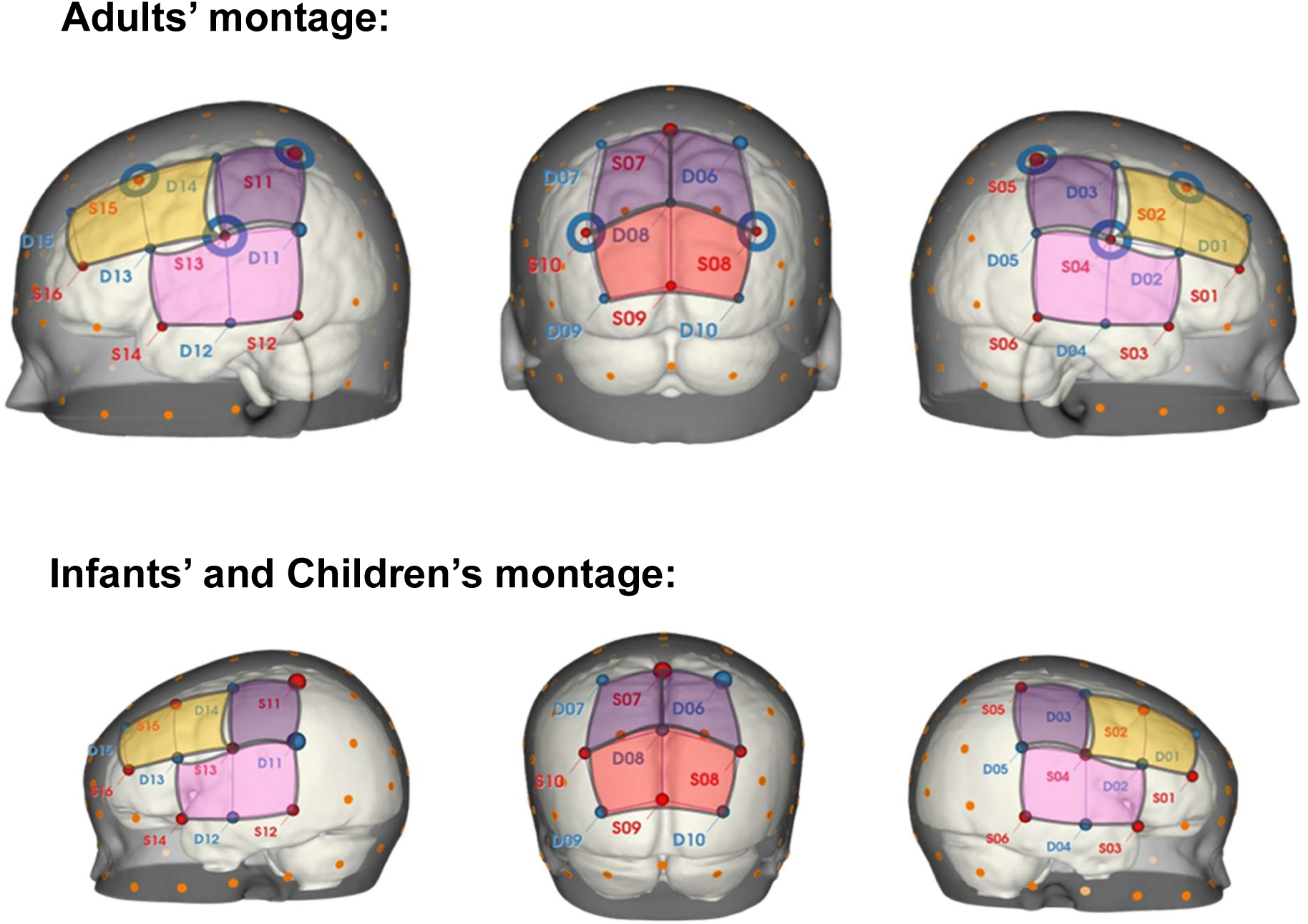
The fNIRS channel montages for adults (top) and children and infants (bottom), from the right (right) posterior (middle) and left (left) views. Detectors are marked in blue, sources are marked in red. Short channels are marked as a blue circle around a source. The approximate ROIs are marked for the frontal (yellow), temporal (pink), parietal (purple) and occipital (red) cortical areas.

#### Data Acquisition

Near-infrared absorption rates at wavelengths of 760 nm and 850 nm were recorded at a sampling rate of 5.08 Hz using Aurora 2021 software (NIRx), with experimental stimuli administered via PsychoPy.

### 2.4 Data Analysis

#### Cap registration

Accurate channel positioning was achieved using the STORM-Net digitizing tool. This tool estimates the channel locations of the fNIRS cap based on a short video of each participant (Erel et al., 2020) that provides a list of MNI coordinates for precise assignment of channels to the ROIs for each participant. The ROIs depicted in Figure 2 are approximate, and channel-to-ROI mapping varied slightly across participants.

#### Preprocessing

The fNIRS signals were preprocessed and analyzed using SATORI software (NIRx). A general linear model (GLM) analysis was conducted with SATORI and further analyzed in MATLAB. Functional connectivity and event-related averages (ERA) used Python’s MNE library, with additional statistical processing in MATLAB and JASP.

For adults, the pre-processing workflow consisted of setting a CV threshold of 10% (Piper et al., 2013), converting RAW wavelength data to Optical Density (OD), and performing spike removal with 10 iterations at a 5s lag, at a threshold of 3.5, an influence of 0.5, and a monotonic interpolation. Temporal filtering was applied with a low-pass Butterworth filter of 0.5 Hz and a high-pass Butterworth filter of 0.01 Hz. Short channel regression was performed using the closest SSR, followed by conversion of the OD data to Concentration (CC) data. The duration of all events was set to 2s in the event manager, and the data were normalized using a Z-transform. Infants’ and children’s pre-processing followed the same workflow but used a CV threshold of 30% (Piper et al., 2013) and omitted the short channel regression.

#### Measuring Adaptation

adaptation was defined as a reduction in HbO amplitude across repeated stimulus presentations within each block. This was measured for each ISI using a GLM (positive beta coefficient indicating full hemodynamic response and no adaptation, negative beta coefficient indicating reduced hemodynamic response and adaptation) and ERA analysis (visually observing the HbO reaction throughout each ROI and ISI, along with anova to examine the differences between age groups).

### 2.5 GLM analysis

The ISIs (stimulus rates) were included in the model as independent variables. A two-gamma HRF model was used, with different HRF parameters for each group as follows:

For the infants, the onset of the response was set at 0 seconds, with the peak response time occurring at 10 seconds. The peak undershoot was 18 seconds. The responses-undershoot ratio was set to 6. The response and undershoot dispersions were both set to 1.

For children, the onset was also set at 0, but the response peak was set at 8 seconds. The time to undershoot peak was reduced to 16 seconds. The response-undershoot ratio remained at 6, and the dispersions of the response and undershoot remained at 1.

For adults, the “default” HRF parameters were used: the onset of the response was set to 0 seconds, the response peak was set to 6 seconds, and the undershoot peak was set to 16 seconds. The response-undershoot ratio remained at 6, and the dispersions of the undershoot and response remained at 1.

### 2.5 Supplementary materials

All stimuli, analysis codes and figures not presented in this paper such as HbR analysis are available on OSF at the following link: https://osf.io/nq2cz/?view_only=e76505040e19485ea436c26243d73cdc

## 3. Results

### 3.1 Memory test

After the fNIRS recording phase, the child and adult participants were presented with a 2AFC identification task where they were asked to indicate which item was presented. Adults scored significantly higher than children on the memory test after the visual task *(Figure 3: 98.26% vs. 85.24% ; p<0.05, t=3.2, df=63)*. Both groups’ responses were above chance.

**Figure 3:**
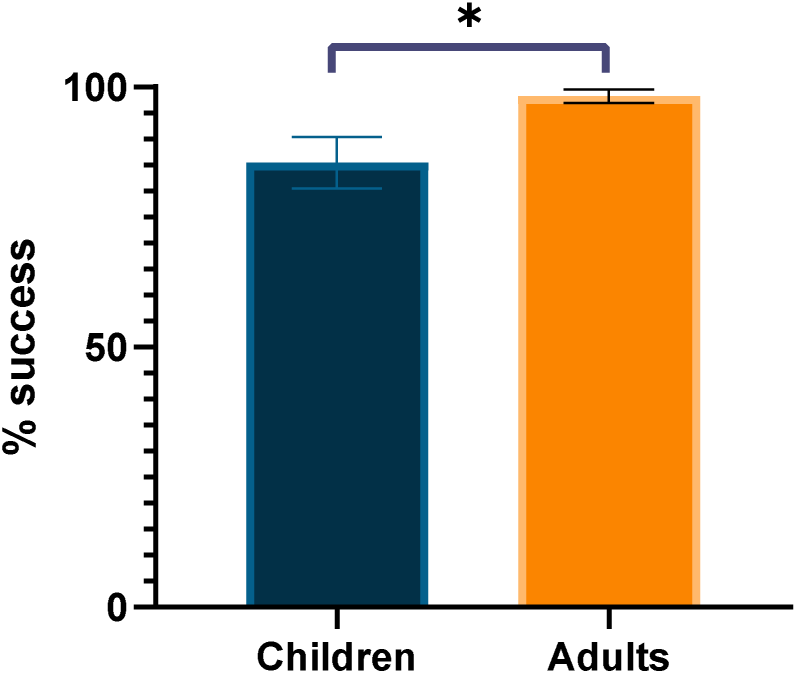
Memory scores for adults and children. Adults (right, orange) showed a significantly higher average success rate than children (left, blue). Error bars denote the standard error of the mean.

### 3.2 Cortical response amplitude age-group effect by ROI

A GLM analysis was employed to quantify the relationship between stimulus presentation rate and neural responses across development. In the GLM analysis, a positive β score indicates a positive hemodynamic response function (HRF), as modeled by the GLM. A main effect between groups was found in the bilateral Frontal *(Figure 4; Right: F(2,87)=11.56, p<0.001, Left: F(2,92)=12.5, p<0.001)*, Temporal *(Right: F(2,93)=12.51, p<0.001, Left: F(2,92)=14.4, p<0.001)*, and the left Parietal *(F(2,93)=3.66, p=0.03)* cortical areas (Figure 4).

**Figure 4:**
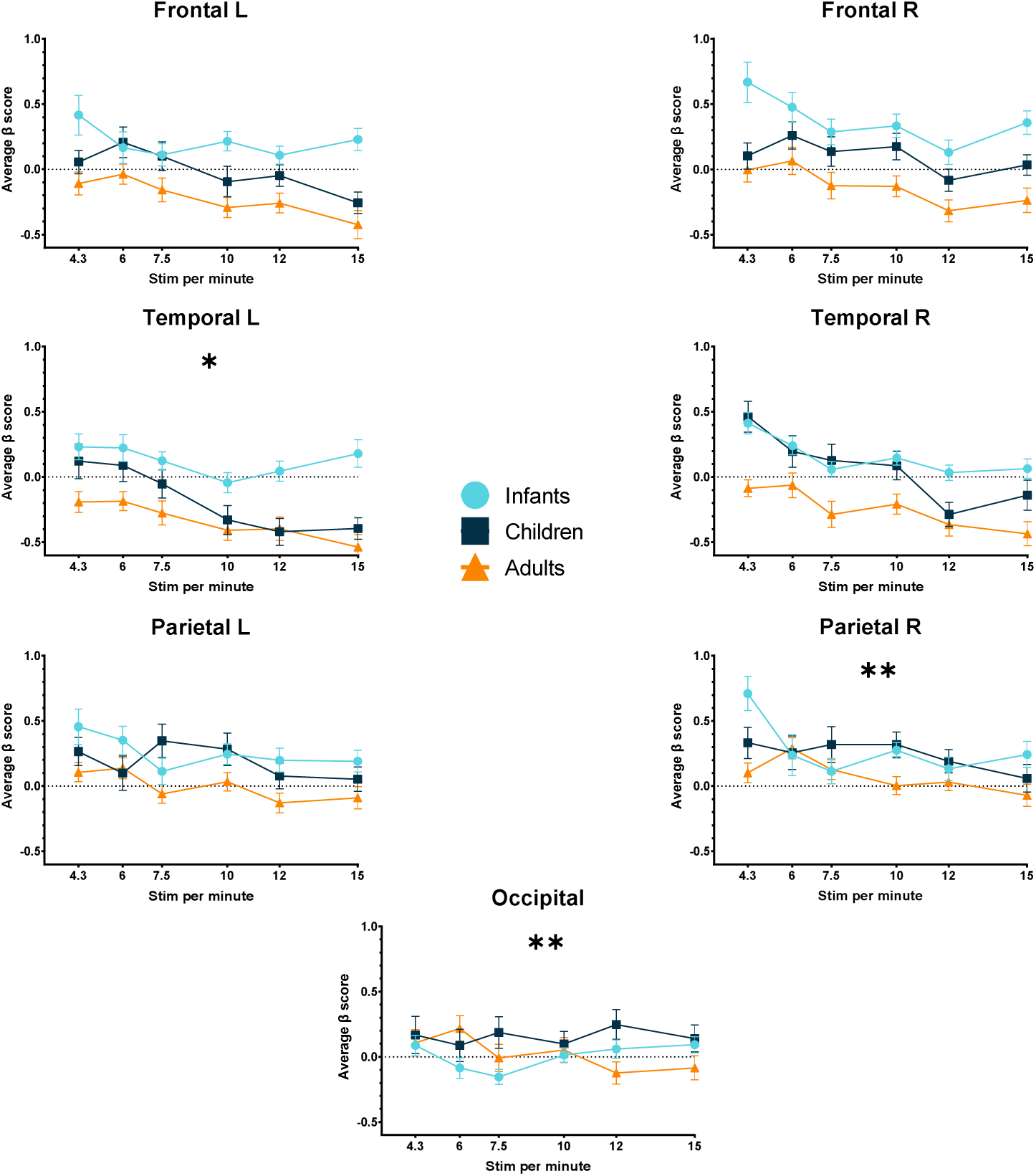
GLM HbO₂ β scores for all ROIs for Infants (light blue), Children (dark blue), and Adults (orange). Average β scores as a function of stimuli per minute by group, where ISIs of 2,3,4,6,8, and12 seconds correspond to 15,12,10,7.5,6, and 4.3 stimuli per minute. A significant age **×** stimulus presentation rate interaction was found for the Left Temporal, Right Parietal, and Occipital ROIs.

For these ROIs, the average β score differed across groups regardless of stimulus rate, with infants having the highest score. This suggests a difference in general brain response to the stimuli in each age group.

A group **×** stimulus rate interaction was found for the Occipital (*F(2,10)=3.18, p<0.001*), Left Temporal (*F(2,10)=1.94, p=0.03*), and Right Parietal (*F(2,10)=3.05, p<0.001*) cortical areas (Table 1). For these ROIs there was a difference in the effect of stimulus presentation rate on brain response (HbO₂), with infants showing faster decay of adaptation than children and adults.

**Table 1:**
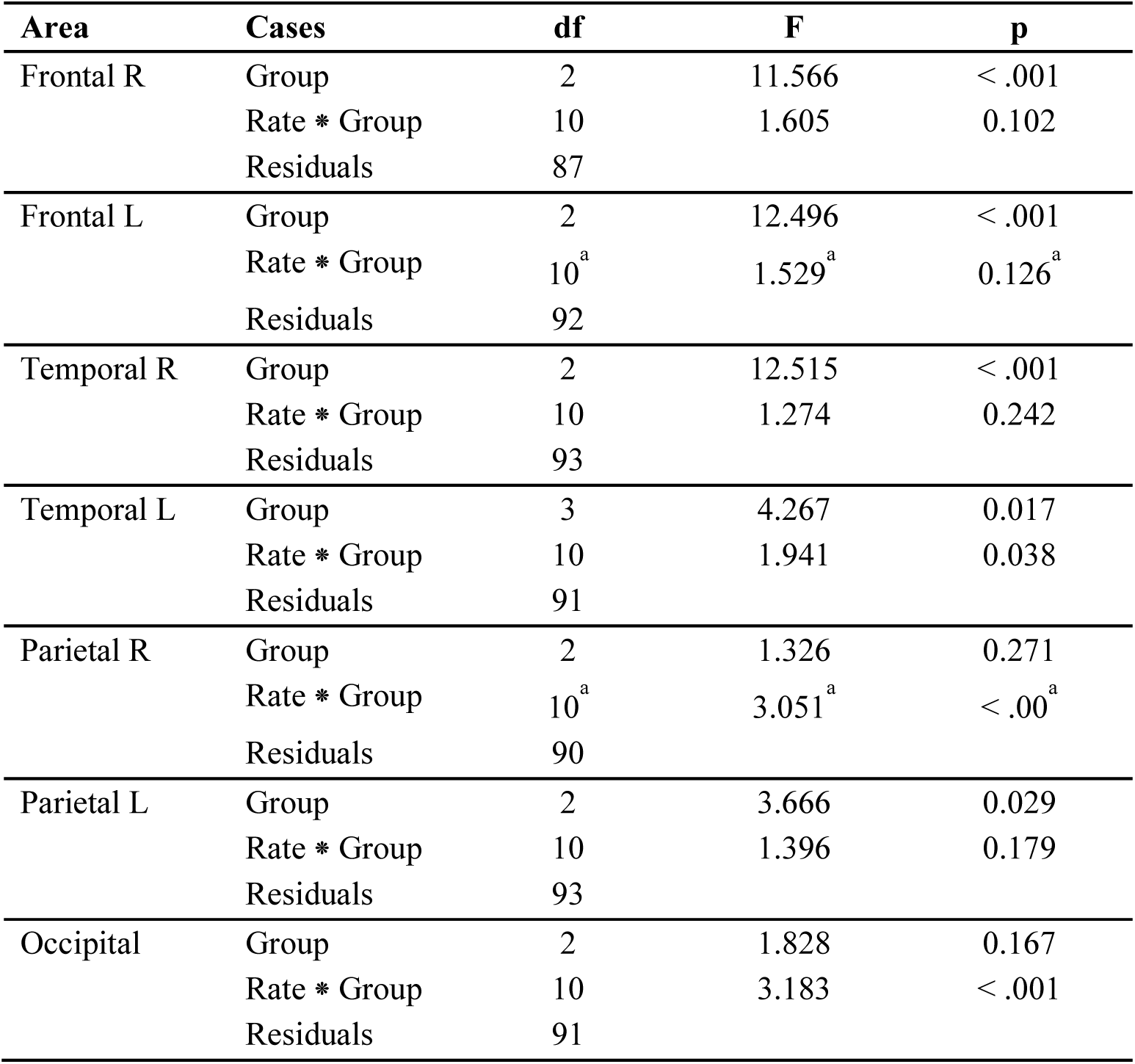
Group effects by ROI. Repeated measures ANOVA, examining the effects of stimulus presentation rate on hemodynamic responses (HbO₂) across ROIs. A main effect was found for the bilateral frontal (*Right: F(2,87)=11.56, p<0.001, Left: F(2,92)=12.5, p<0.001*) and temporal (*Right: F(2,93)=12.51, p<0.001, Left: F(2,92)=14.4, p<0.001*) cortex areas, as well as the left parietal cortex (*F(2,93)=3.66, p=0.03*). A significant rate x group interaction was found for the left temporal (*F(2,10)=1.94, p=0.03*), right parietal (*F(2,10)=3.05, p<0.001*) and occipital (*F(2,10)=3.18, p<0.001*) ROIs.

| Area | Cases | df | F | p |
| --- | --- | --- | --- | --- |
| Frontal R | Group | 2 | 11.566 | < .001 |
|  | Rate * Group | 10 | 1.605 | 0.102 |
|  | Residuals | 87 |  |  |
| Frontal L | Group | 2 | 12.496 | < .001 |
|  | Rate * Group | 10<sup>a</sup> | 1.529<sup>a</sup> | 0.126<sup>a</sup> |
|  | Residuals | 92 |  |  |
| Temporal R | Group | 2 | 12.515 | < .001 |
|  | Rate * Group | 10 | 1.274 | 0.242 |
|  | Residuals | 93 |  |  |
| Temporal L | Group | 3 | 4.267 | 0.017 |
|  | Rate * Group | 10 | 1.941 | 0.038 |
|  | Residuals | 91 |  |  |
| Parietal R | Group | 2 | 1.326 | 0.271 |
|  | Rate * Group | 10<sup>a</sup> | 3.051<sup>a</sup> | < .00<sup>a</sup> |
|  | Residuals | 90 |  |  |
| Parietal L | Group | 2 | 3.666 | 0.029 |
|  | Rate * Group | 10 | 1.396 | 0.179 |
|  | Residuals | 93 |  |  |
| Occipital | Group | 2 | 1.828 | 0.167 |
|  | Rate * Group | 10 | 3.183 | < .001 |
|  | Residuals | 91 |  |  |

**Table 2:** Multiple comparisons Anova group effects of functional connectivity. Several ROI pairings show statistically significant age-related differences (p_FDR< 0.05), with particularly strong effects observed in Temporal R*Temporal L, Parietal R*Temporal R and Parietal R*Temporal L (p_FDR< 0.0001). Several pairings showed no significant difference between the age groups such as the bilateral Frontal, Temporal R*Occipital, Parietal R*Frontal L, Frontal L*Occipital, Temporal L*Occipital (p_FDR >0.05).

| ROI pairings | Df2 | F | p_FDR | $\eta^2p$ |
| --- | --- | --- | --- | --- |
| Frontal R* Temporal R | 97 | 4.84 | 0.01* | 0.091 |
| Frontal R* Parietal R | 97 | 6.29 | 0.006** | 0.115 |
| Frontal R* Frontal L | 97 | 0.97 | 0.40 | 0.020 |
| Frontal R* Temporal L | 96 | 7.60 | 0.002** | 0.137 |
| Frontal R* Parietal L | 97 | 3.62 | 0.043* | 0.069 |
| Frontal R* Occipital | 89 | 5.20 | 0.013* | 0.105 |
| Temporal R * Parietal R | 97 | 14.79 | < .0001**** | 0.234 |
| Temporal R * Frontal L | 99 | 4.65 | 0.019* | 0.086 |
| Temporal R * Temporal L | 99 | 12.94 | < .0001**** | 0.207 |
| Temporal R * Parietal L | 98 | 5.91 | 0.008** | 0.108 |
| Temporal R * Occipital | 89 | 0.96 | 0.40 | 0.021 |
| Parietal R * Frontal L | 97 | 1.97 | 0.16 | 0.039 |
| Parietal R * Temporal L | 96 | 11.92 | 0.00012*** | 0.199 |
| Parietal R * Parietal L | 97 | 3.50 | 0.044* | 0.067 |
| Parietal R * Occipital | 89 | 11.48 | 0.0001*** | 0.205 |
| Frontal L * Temporal L | 98 | 7.09 | 0.003** | 0.126 |
| Frontal L * Parietal L | 98 | 4.31 | 0.024* | 0.081 |
| Frontal L * Occipital | 89 | 2.95 | 0.07 | 0.062 |
| Temporal L * Parietal L | 98 | 14.05 | < .0001**** | 0.223 |
| Temporal L * Occipital | 89 | 0.42 | 0.65 | 0.009 |
| Occipital * Parietal L | 90 | 9.87 | 0.0004*** | 0.180 |

*Note:* Df1 equals 2

### 3.3 Full block brain response analysis - Event Related Averages (ERA)

To complement the GLM findings and visualize the temporal dynamics of the neural responses, we calculated Event-Related Averages (ERAs). ERA provides a more detailed view of how the hemodynamic response evolves over time, and thus can reveal adaptation patterns that might not be fully captured by parametric models. This visualization indicated adaptation in the lower ISIs (higher rates of stimulus presentation), particularly in the adult group: the brain response decreased until reaching a plateau (asymptote) and returned to baseline after the block ended.

This pattern aligns with typical adaptation brain responses (Arichi et al., 2012). However, at higher ISIs (lower rates of stimulus presentation), children and infants showed stronger brain responses with identical responses to each stimulus, which is consistent with recovery from adaptation. Overall, the groups displayed the greatest differences in brain responses for shorter ISIs but exhibited greater similarities for longer ISIs.

#### 3.3.1 Frontal stimulus evoked response by ISI

For the shortest ISIs, the adults showed strong adaptation with a marked reduction in response, whereas children and infants displayed signs of recovery throughout the block (Figure 5). Adults maintained this adaptation pattern up to the longest ISIs, where their response, although no longer showing typical adaptation, still did not show full recovery. By contrast, infants and children showed recovery from adaptation at the 6-second ISI and achieved full recovery at the longest 12-second ISI.

**Figure 5:**
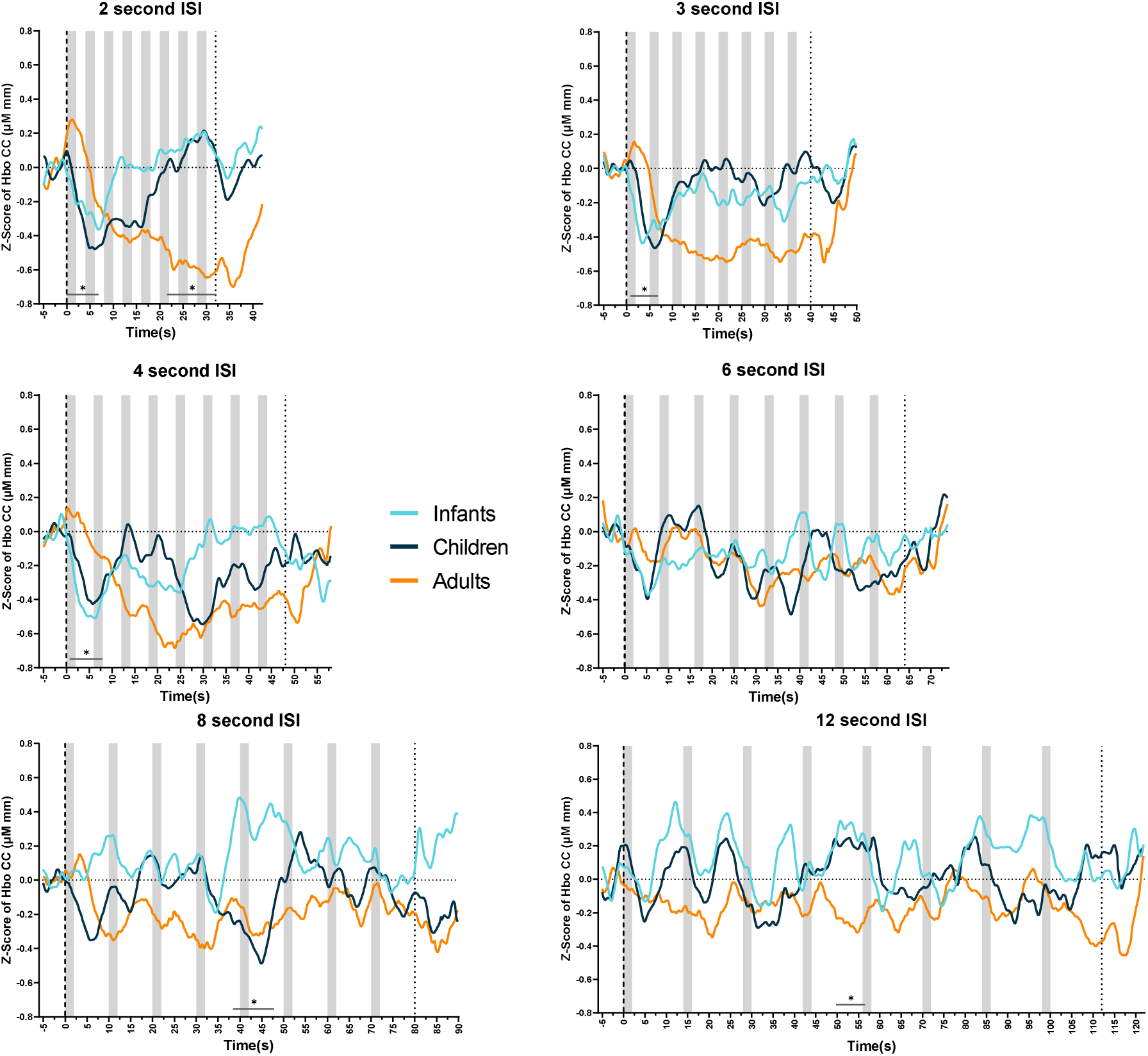
Average brain response in the bilateral frontal cortex for infants (light blue), children (dark blue) and adults (orange). Z-score of HbO₂ concentration measured starting 5 seconds before the first stimulus to 10 seconds after the end of each block. Gray rectangles mark each stimulus presentation. Black bars with * mark timepoints where the differences between groups were significant (p<0.5) **A.** 2-second ISI (15 stimuli per minute) **B.** 3-second ISI **C.** 4-second ISI **D.** 6-second ISI **E.** 8-second ISI **F.** 12-second ISI.

#### 3.3.2 Temporal stimulus evoked response by ISI

At a 2-second ISI, adults and children showed strong adaptation, whereas infants did not display typical adaptation. At 3-, 4-, and 6-second ISIs, all groups demonstrated adaptation to the stimulus (Figure 6). At 8- and 12-second ISIs, infants exhibited a strong response resembling recovery from adaptation. Adults and children had not yet fully recovered, with children showing a reduced response even at the highest ISI.

**Figure 6:**
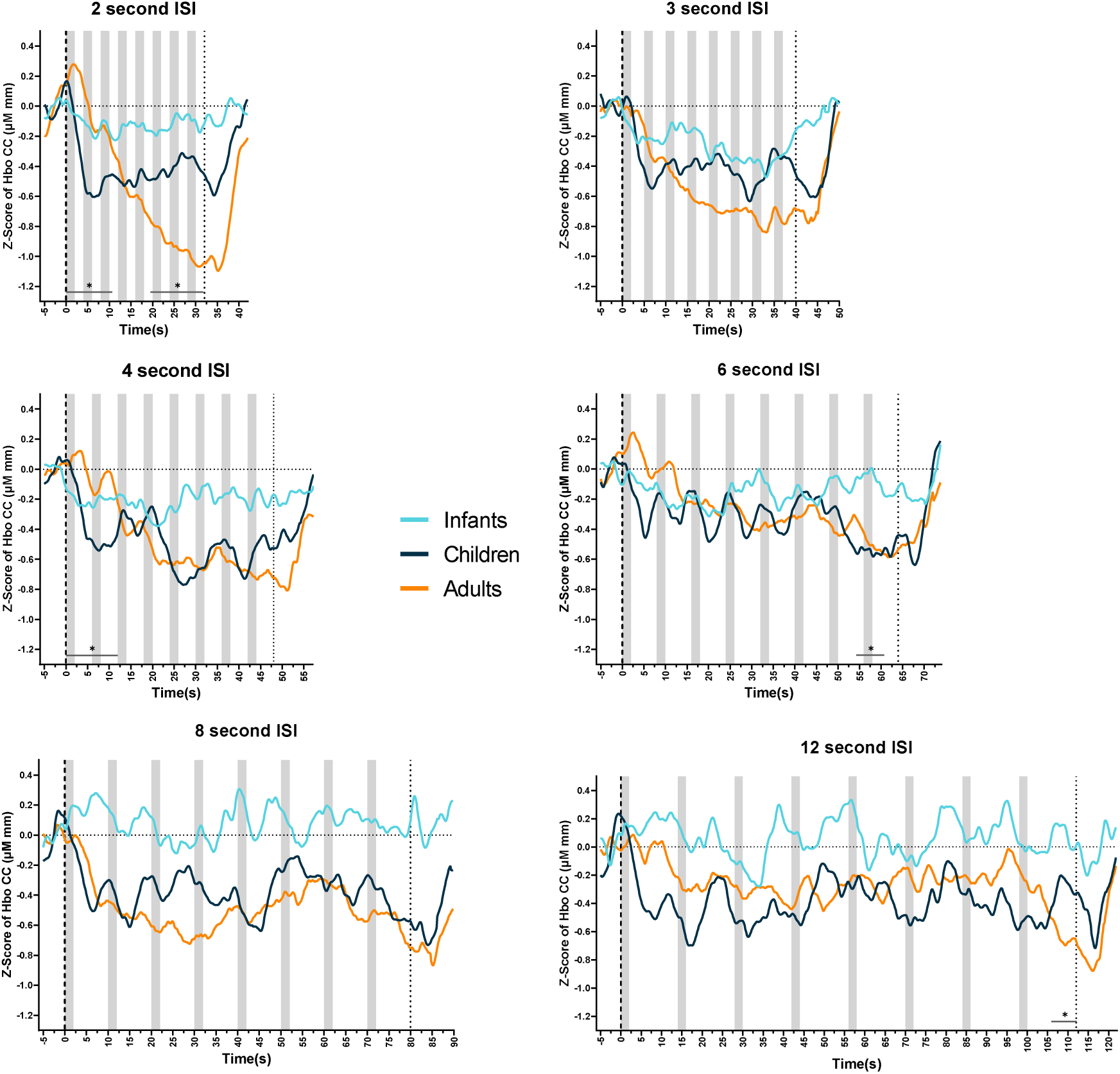
Average brain response in the bilateral temporal cortex for infants (light blue), children (dark blue) and adults (orange). Z-score of HbO₂ concentration measured starting 5 seconds before the first stimulus to 10 seconds after the end of each block. Gray rectangles mark each stimulus presentation. Black bars with * mark timepoints where the differences between the groups were significant (p<0.5) **A.** 2-second ISI (15 stimuli per minute) **B.** 3-second ISI **C.** 4-second ISI **D.** 6-second ISI **E.** 8-second ISI **F.** 12-second ISI.

#### 3.3.3 Parietal stimulus evoked response by ISI

At 2-, 3-, and 4-second ISIs, adults showed strong adaptation to the stimulus, whereas infants displayed recovery for these same intervals (Figure 7). The children’s responses were comparable to infants at the lowest ISI but showed robust brain activity at 3- and 4-second ISIs. At the 6-second ISI, infants demonstrated strong adaptation, while adults and children exhibited recovery. At the highest ISIs of 8 and 12 seconds, all groups showed strong responses after each stimulus presentation, consistent with recovery from adaptation.

**Figure 7:**
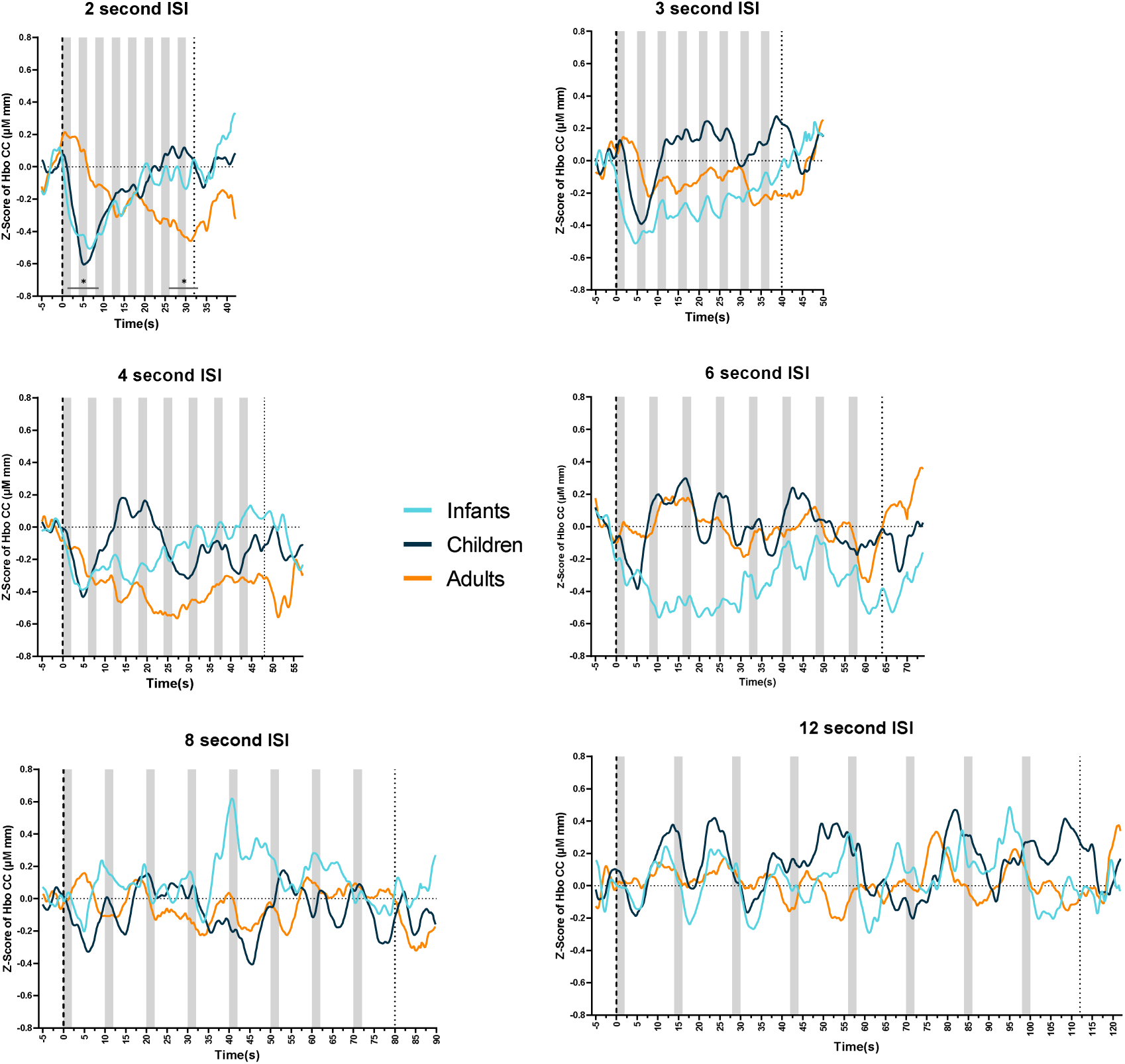
Average brain response in the bilateral parietal cortex for infants (light blue), children (dark blue) and adults (orange). Z-score of HbO₂ concentration measured starting 5 seconds before the first stimulus to 10 seconds after the end of each block. Gray rectangles mark each stimulus presentation. Black bars with * mark timepoints where the differences between the groups were significant (p<0.5) **A.** 2-second ISI (15 stimuli per minute) **B.** 3-second ISI **C.** 4-second ISI **D.** 6-second ISI **E.** 8-second ISI **F.** 12-second ISI.

### 3.4 Functional Connectivity

The functional connectivity analysis made it possible to assess how the brain regions coordinated activity during sensory processing, thus reflecting the maturation of neural networks across development. By examining correlation patterns between the different cortical areas, we could evaluate whether age-related differences in adaptation were paralleled by corresponding changes in network integration. This approach provided critical insights into whether the developmental trajectory of sensory processing reflects isolated regional changes or broader network reorganization.

#### 3.4.1 Correlation patterns for each age-group

We computed mean Pearson correlation coefficients (r) between ROI pairs for each participant and then averaged these coefficients across participants to generate group-level connectivity matrices (Figure 8A). Each cell in the matrix shows the Pearson’s correlation coefficient (r) between the activity patterns of two ROIs during the experiment. The analysis revealed distinct patterns across age groups as follows:

**Figure 8:**
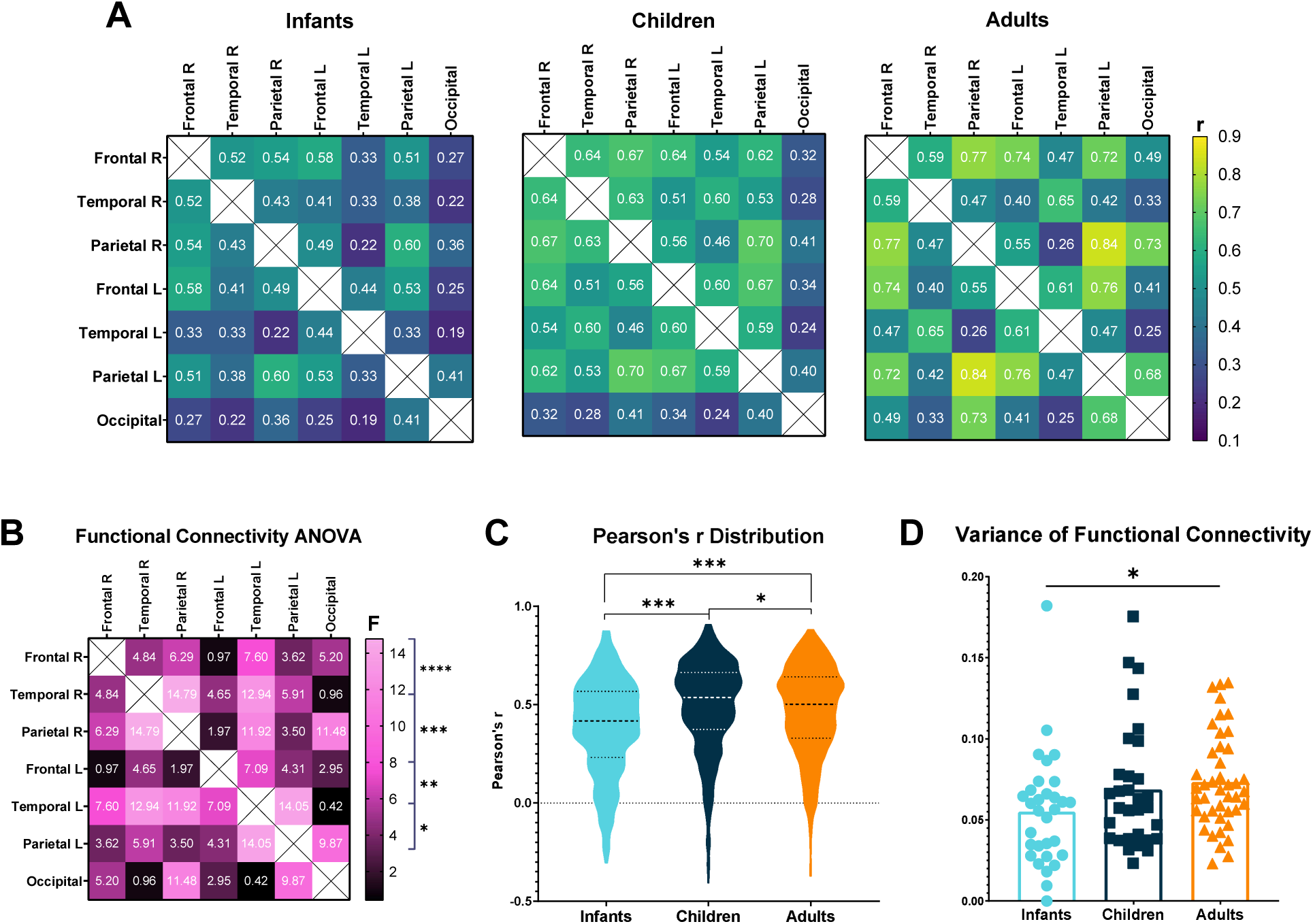
**A:** Functional Connectivity Correlation Matrix. The x and y axes show brain regions (ROIs): Frontal R/L, Temporal R/L, Parietal R/L, and Occipital. Correlation coefficient (r) indicates the strength of the functional connectivity between ROI pairs. Values closer to 1 indicate stronger connectivity. **B:** Functional connectivity matrix comparing the F values of the repeated measures ANOVA for each ROI pairing. Higher F scores are pinker whereas lower F scores are blacker. The p-value significance (\**p<0.05*, *\*\*p<0.01, ***p<0.001*, etc.) corresponds to the right color bar. **C:** Distribution of infants’ (blue), children’s (red) and adults’ (green) individual Pearson’s R correlation values for all ROIs. The y axis shows the Pearson’s r value. **D:** Distribution of the variances for infants (blue), children (dark blue) and adults (orange).

Adults demonstrated the strongest connectivity between the bilateral parietal regions (r=0.84) and the right frontal-parietal regions *(r=0.77)*, but weaker connections between the left frontal-right parietal (r=0.26) and the left temporal-occipital regions *(r=0.25)*. Children showed high connectivity across most ROIs, except for the occipital cortex. Their strongest connections paralleled adults’: the bilateral parietal *(r=0.7)* and right frontal-parietal regions *(r=0.67)*. The temporal-occipital connections were weaker, mirroring adults *(left: r=0.24, right: r=0.28)*. Infants exhibited lower overall connectivity, possibly reflecting ongoing development or measurement noise. Their strongest connections were between the bilateral parietal *(r=0.6)* and the bilateral frontal regions *(r=0.58)*, with weaker temporal-occipital (*left: r=0.19, right: r=0.22)* and right parietal-left temporal *(r=0.22)* connections.

Further visual inspection of the functional connectivity matrices revealed developmental differences in network organization. The adult FC matrix appeared more "mountainous", with pronounced peaks and valleys reflecting strong or weak connectivity between regions. In contrast, the infant FC matrix was “flatter," indicating lower overall connectivity and a more uniform distribution of correlations. This pattern may reflect the immaturity of large-scale network integration and suggests that neural connectivity becomes more specialized with age.

#### 3.4.2 ROI specific functional connectivity

ANOVA analysis compared functional connectivity differences across age groups, with FDR correction for multiple comparisons (Benjamini & Hochberg, 1995). Results revealed significant age-related differences in functional connectivity between multiple brain regions (Figure 8B). Post hoc analyses revealed that the most significant differences occurred between infants and adults, though the specific pattern of connectivity changes varied across ROI pairs. Notably, bilateral frontal connections (consistently high) and temporal-occipital connections (consistently low) showed no significant group differences, suggesting these networks establish their connectivity early in development while others undergo maturation.

#### 3.4.3 Overall functional connectivity

Examining the overall distribution of functional connectivity values across all brain regions provided a holistic view of neural network integration across development (figure 8C).

Significant differences were found between age groups when looking at the distributions of all Pearson’s r correlations of each group, regardless of ROI *(F(2,84)=5.65, p=0.005).* These fundamental differences in cortical connectivity patterns may underlie the varying adaptation responses observed between age groups, thus providing insights into how neural maturation affects broad information processing capabilities.

A variance analysis of the arctanh of each subject (Figure 8D) indicated a significant difference between groups (p=0.023). This reflects the different “terrains” of each age group’s functional connectivity matrices.

A post-hoc analysis of the repeated measures ANOVA for the functional connectivity (Table 3) revealed a significantly lower average correlation value for infants compared to children *(t=-10.31, p<0.001)* and adults *(t=-8.54, p<0.001)*, and a significantly higher average correlation value for the children compared to adults *(t=2.62, p=0.02)*.

**Table 3:** Post hoc comparisons of repeated measures ANOVA for the functional connectivity analysis. Both Tukey and Bonferroni tests were used for the analysis.

| Group 1 | Group 2 | Mean difference | SE | t | ptukey | pbonf |
| --- | --- | --- | --- | --- | --- | --- |
| Infants | Children | -0.159 | 0.015 | -10.315 | < .001 | < .001 |
|  | Adults | -0.124 | 0.014 | -8.547 | < .001 | < .001 |
| Children | Adults | 0.035 | 0.013 | 2.629 | 0.023 | 0.026 |

*Note.* P-value adjusted for comparing a family of 3
*Note.* Results are averaged over the levels of: ROIs

## 4. Discussion

### 4.1 Aims and Hypotheses

This study investigated the developmental trajectory of cortical adaptation across infants, children, and adults using functional near-infrared spectroscopy (fNIRS). The primary objectives were to determine whether cortical adaptation recovery varies across age groups and to characterize developmental changes in functional connectivity patterns between cortical regions.

To examine these questions, participants across three developmental stages were presented with auditory-visual stimuli at varying inter-stimulus intervals (ISIs), while hemodynamic responses were measured in the frontal, parietal, temporal, and occipital cortical areas. We hypothesized that infants would demonstrate more rapid recovery from adaptation than children and adults, and that adults would exhibit the slowest recovery. We also predicted that functional connectivity would be the weakest in infants and progressively stronger in children and adults.

### 4.2 Explicit Memory Development

As expected, adults significantly outperformed children on explicit memory tests, consistent with well-confirmed findings demonstrating developmental improvements in working memory capacity from early childhood through adulthood (Bauer et al., 2017).

### 4.3 Age-Related Differences in Cortical Adaptation

#### 4.3.1 GLM Analysis: Developmental Progression in Adaptation Recovery

The General Linear Model (GLM) analysis revealed significant main effects for age in the bilateral frontal, bilateral temporal, and left parietal regions of interest (ROIs). While all groups showed decreased brain responses with increased stimulus presentation rates, the magnitude of this effect varied substantially across age groups.

In particular, there were significant age × ISI interactions in the occipital, left temporal, and right parietal ROIs. The differential impact of ISI on hemodynamic responses across age groups in these primarily sensory regions suggests fundamental developmental differences in sensory processing mechanisms. Consistent with our hypothesis, infants had higher brain responses to longer ISIs (lower stimulus presentation rates) compared to adults, with children showing intermediate responses.

#### 4.3.2 ERA Analysis: Age-Dependent Adaptation Dynamics

The ERA graphs provide a more comprehensive visualization of the hemodynamic responses across the entire experimental paradigm compared to the GLM analysis. Adults exhibited clear adaptation at shorter ISIs with minimal recovery, whereas infants demonstrated strong recovery across longer ISIs, in line with our initial hypothesis. Although no clearcut ISI threshold for adaptation recovery emerged for any group or ROI, all groups showed some degree of recovery at the longest ISI.

It is worth noting that the observed hemodynamic responses were generally lower in amplitude than typically reported for these age groups (Arichi et al., 2012; Lloyd-Fox et al., 2019; Moses et al., 2013; Richter & Richter, 2003). This may be attributable to the brief stimulus duration (2 seconds), which might have been insufficient to elicit full hemodynamic responses. This could also explain why the GLM analysis failed to detect recovery from adaptation in adults for the longest ISI, unlike the ERA visualization.

### 4.4 Functional Connectivity Patterns across Development

The strongest correlations across all age groups were observed between parallel regions: the left and right parietal areas and left and right frontal cortical areas. Both children and adults exhibited robust inter-ROI correlations, whereas infants displayed significantly weaker inter-ROI correlations and lower average correlation values. This pronounced difference in connectivity patterns may reflect the neural immaturity characteristic of early development (Emberson et al., 2015; Gao et al., 2014; Gualtieri & Finn, 2022).

### 4.5 Conclusions and Implications

The multifaceted analysis presented in this study provides valuable insights into cortical adaptation across development. The GLM analysis revealed significant age × ISI interactions in the right parietal, left temporal, and occipital ROIs. The ERA analysis further elucidated these interactions by demonstrating age-dependent adaptation dynamics: infants exhibited rapid recovery from adaptation, particularly at longer ISIs, adults demonstrated slower recovery, which was only evident at the longest ISI, and children’s responses were intermediate. The functional connectivity analysis revealed a developmental progression, with adults and children showing similar patterns of strong inter-ROI correlations, particularly between the bilateral frontal and parietal regions, whereas infants displayed significantly lower overall inter-ROI correlations.

These results have major implications for our understanding of neural development and sensory processing. The observed age-related differences in cortical adaptation suggest that the brain’s ability to filter and process repeated stimuli evolves throughout development. The stronger responses by infants to longer stimulus intervals, combined with their lower inter-ROI correlations, may reflect a neural system optimized for maximal information intake and learning, albeit with less efficient filtering of redundant information. This aligns with theories of early brain development that emphasize heightened plasticity and learning capacity in infancy (Kolb et al., 2017; Werker & Hensch, 2014). The similarities in functional connectivity between children and adults, despite differences in adaptation patterns, suggest that the basic architecture for sensory processing is established relatively early, whereas the fine-tuning of adaptive responses continues to develop (Cai et al., 2018; Homae et al., 2010). These findings may help explain previously observed behavioral differences across age groups, such as the superior performance of children in tasks requiring "in-the-moment" perception versus adults’ greater reliance on prior experience (Jaffe-Dax et al., 2023).

Collectively, these findings suggest that cortical adaptation mechanisms undergo substantial developmental changes, with implications for how the brain processes and integrates sensory information across the lifespan.

### 4.6 Limitations

Several methodological limitations warrant consideration. While GLM analysis is a standard approach for fNIRS data, it presents certain challenges in developmental research. No standardized 2- Gamma HRF model exists for infants and children, necessitating either the application of adult models or, as in our study, the development of group-specific models based on observed responses. This approach potentially introduced bias by using the same data for model construction and analysis. Although we employed complementary methods such as functional connectivity analysis, whole-block visualization, and ERA analysis to validate and compare results, future studies should aim to develop more reliable, age-appropriate HRF models to enhance the accuracy and reliability of GLM analysis in developmental research.

While the use of unfamiliar abstract stimuli allowed for controlled comparisons between age groups, it may not have reflected how the brain processes more ecologically valid stimuli encountered in daily life (Emberson et al., 2016; Lloyd-Fox et al., 2019). Furthermore, while fNIRS offers numerous advantages for developmental research, it cannot capture adaptation patterns in subcortical regions that may play crucial roles in sensory processing (Quaresima & Ferrari, 2019).

### 4.7 Future Directions

Future research should address both the limitations of this study and expand the state of the art on cortical adaptation across development. One critical direction would be to generate reliable HRF models specifically for children and infants. Studies should also investigate adaptation using varied stimulus durations and more naturalistic stimuli to better reflect real-world contexts. Further, longitudinal research would offer advantages over our cross-sectional approach by mapping a more precise developmental trajectory of cortical adaptation.

Extending previous work by Jaffe-Dax et al. (2017, 2018), future studies should include clinical populations and explore cortical adaptation in neurodevelopmental disorders such as autism spectrum disorder (ASD) and dyslexia. This approach could identify whether sensory processing differences in these conditions emerge early in development or evolve over time.

In general, future work can build on the foundation established by this study to develop a more comprehensive understanding of how cortical adaptation develops and functions throughout life. This will not only advance theoretical understanding of brain development but may also inform interventions for individuals with atypical sensory processing.

